# Fragmentation of pooled PCR products for highly multiplexed TILLING

**DOI:** 10.1101/451567

**Authors:** Andrea Tramontano, Luka Jarc, Joanna Jankowicz-Cieslak, Bernhard J. Hofinger, Katarzyna Gajek, Miriam Szurman-Zubrzycka, Iwona Szarejko, Ivan Inglebrecht, Bradley J. Till

## Abstract

Improvements to massively parallel sequencing have allowed the routine recovery of natural and induced sequence variants. A broad range of biological disciplines have benefited from this, ranging from plant breeding to cancer research. The need for high sequence coverage to accurately recover single nucleotide variants and small insertions and deletions limits the applicability of whole genome approaches. This is especially true in organisms with a large genome size or for applications requiring the screening of thousands of individuals, such as the reverse-genetic technique known as TILLING, where whole genome approaches become cost prohibitive. Using PCR to target and sequence chosen genomic regions provides an attractive alternative as the vast reduction in interrogated bases means that sample size can be dramatically increased through amplicon multiplexing and multidimensional sample pooling while maintaining suitable coverage for recovery of small mutations. Direct sequencing of PCR products is limited, however, due to limitations in read lengths of many next generation sequencers. In the present study we show the optimization and use of ultrasonication for the simultaneous fragmentation of multiplexed PCR amplicons for TILLING highly pooled samples. Thirty-two PCR products were produced from genomic DNA pools representing 265 pooled barley mutant lines. Mutant lines were produced with the chemical mutagen ethyl methanesulfonate. Samples were subjected to 2×300PE illumina sequencing. Evaluation of read coverage and base quality across amplicons suggests this approach is suitable for high-throughput TILLING and other applications employing highly pooled complex sampling schemes. Induced mutations previously identified in a traditional TILLING screen were recovered in this dataset further supporting the efficacy of the approach.

## Background

Next generation sequencing (NGS) techniques have had a profound impact on biological research. While technologies continue to advance, whole genome approaches remain costly for projects involving the analysis of species with large genomes or those involving the interrogation of many individuals. A variety of reduced representation genome sequencing approaches have been described to circumvent this issue ^1^. One powerful approach to evaluate sequence variation in targeted regions is by sequencing PCR amplicons. Amplicon sequencing has been applied for the discovery and characterization of both natural and induced mutations in plant and animal populations ^2–5^. For example, sequencing has been used to increase throughput of mutation discovery for the reverse-genetics technique known as TILLING (Targeting Induced Local Lesions IN Genomes)^6^,^7^. For TILLING, mutant populations are typically created using mutagens such as ethyl methanesulfonate (EMS) that induce primarily single nucleotide variants. While the frequency of mutations varies between species and ploidy level, mutation densities in diploid species have been reported to range between 1 mutation per 150,000 and 1 mutation per 700,000 base pairs^8^. TILLING screens typically aim to recover multiple mutations from a single target to increase the chances of recovering deleterious alleles. Therefore, a typical TILLING assay involves the screening of 3000 or more mutant individuals. To increase throughput, genomic DNAs are pooled together prior to mutation screening. The application of new sequencing technologies to improve TILLING throughput has been termed TILLING by Sequencing ^3^. In addition to pooling genomic DNA, multiple amplicons are produced from each gDNA pool and combined prior to massively parallel sequencing to further improve throughput. The quantitative nature of NGS methods means that rare induced and natural SNV mutations can be effectively recovered from pools of over 200 individuals ^4,5^. The most common sequencing platform used for TILLING by Sequencing is Illumina ^3,4,9^. Similar approaches have been taken in population genetics studies to identify rare alleles in large germplasm collections^5^.

Direct sequencing of PCR amplicons is advantageous in that an even read coverage across the target is ensured, thus providing consistent recovery of sequence variants. This approach is disadvantageous, however, due to the fact that the length of sequenced amplicons is limited to the maximum read lengths of the sequencer used (e.g., 500-600 bp for the Illumina MiSeq)^4,5^. Thus, multiple amplicons are required to screen an entire gene, necessitating extra liquid handling steps and also additional work in adjusting PCR amplicon concentrations to a similar level prior to sequencing. An alternative approach is to produce longer PCR products and to fragment them prior to library construction. Several approaches for fragmentation have been described such as nebulisation, enzymatic cleavage, and ultrasonication^10^. Most approaches are developed for the fragmentation of genomic DNA samples that contain high molecular weight molecules. We describe here the optimization of ultrasonication for PCR products amplified from the genomes of *Coffea arabica* (Arabica coffee) and *Hordeum vulgare* (barley) ranging between 670 and 1513 bp. We further show suitable coverage can be achieved in a pool of 32 distinct PCR products generated from PCR amplification of pools of 256 genomic DNA samples prepared from chemically mutagenized barley. This suggests that the approach can be easily adapted for pooled amplicon sequencing in different species.

## Results

### Evaluation of sonication parameters established for genomic DNA on PCR products

A PCR product of 964 base pairs (bp) amplified from *Coffea arabica* genomic DNA was created for initial experiments to optimize DNA fragmentation. The aim of the work was to identify conditions resulting in a fragmented PCR product between 350 and 500 bp using a Corvaris focused-ultrasonicator. We initially tested the manufacturer’s parameters established for genomic DNA shearing (see Supplementary Table S1). Fragment analysis showed limited fragmentation and retention of a high concentration of 964 bp product at all pre-defined parameters (see Supplementary Fig. S1).

### Optimization of parameters for sonication of a single PCR amplicon

Based on the results from the default parameters for genomic DNA we hypothesized that halving the power, duty factor and/or cycles/burst may improve fragmentation of lower molecular weight PCR products. Fourteen distinct sonication parameter combinations were chosen and evaluated (Table 1). Test numbers 1 through 3 were designed to assess the effect of the number of cycles/burst on the fragment size and distribution. Tests 4 through 6 were designed to assess the effect of the duty factor on the fragment size and distribution. Tests 7 through 9 were designed to assess the effect of the peak power on the fragment size and distribution. Tests 10 through 12 were designed to assess the effect of the duration of the sonication on the fragment size and distribution, and tests 13 and 14 were designed to test the combination of high duty factor with an average number of cycles/burst and very low duty factor with the highest cycles/burst respectively.

**Table 1:** Test parameters for sonication of a 964 bp PCR product.

| Test # | Time [s] | Peak power<br>[W] | Duty Factor<br>[%] | Cycles/burst | Volume<br>[μl] | Average power<br>[W] |
| --- | --- | --- | --- | --- | --- | --- |
| 1 | 30 | 75 | 20 | 50 | 60 | 15 |
| 2 | 30 | 75 | 20 | 350 | 60 | 15 |
| 3 | 30 | 75 | 20 | 750 | 60 | 15 |
| 4 | 30 | 75 | 5 | 200 | 60 | 3.75 |
| 5 | 30 | 75 | 25 | 200 | 60 | 18.75 |
| 6 | 30 | 75 | 12,5 | 200 | 60 | 9.38 |
| 7 | 30 | 20 | 50 | 200 | 60 | 10 |
| 8 | 30 | 30 | 50 | 200 | 60 | 15 |
| 9 | 30 | 40 | 50 | 200 | 60 | 20 |
| 10 | 15 | 75 | 20 | 200 | 60 | 15 |
| 11 | 45 | 75 | 20 | 200 | 60 | 15 |
| 12 | 60 | 75 | 20 | 200 | 60 | 15 |
| 13 | 30 | 50 | 40 | 200 | 60 | 20 |
| 14 | 30 | 75 | 5 | 1000 | 60 | 3.75 |

The extent of PCR fragmentation was evaluated using a capillary based Fragment Analyzer (FA), (see Supplementary Fig. S2). We chose to qualitatively evaluate the relative concentration of DNA fragments owing to the fact that the absolute concentration varies according to amount of input PCR product used in the assay. Based on this analysis we concluded that parameters of test # 13 provided the best fragmentation as it produced a broad distribution of fragments with a peak of 390bp, which is in the size range of 350 to 550 bp recommended for Illumina library preparation (Fig. 1)^11^. Additional modifications whereby average power (Peak Power and Duty Factor) were held constant while time, cycles/ burst and volume were modified showed no substantial improvement (see Supplementary Table S2 and Supplementary Fig. S3).

**Figure 1:**
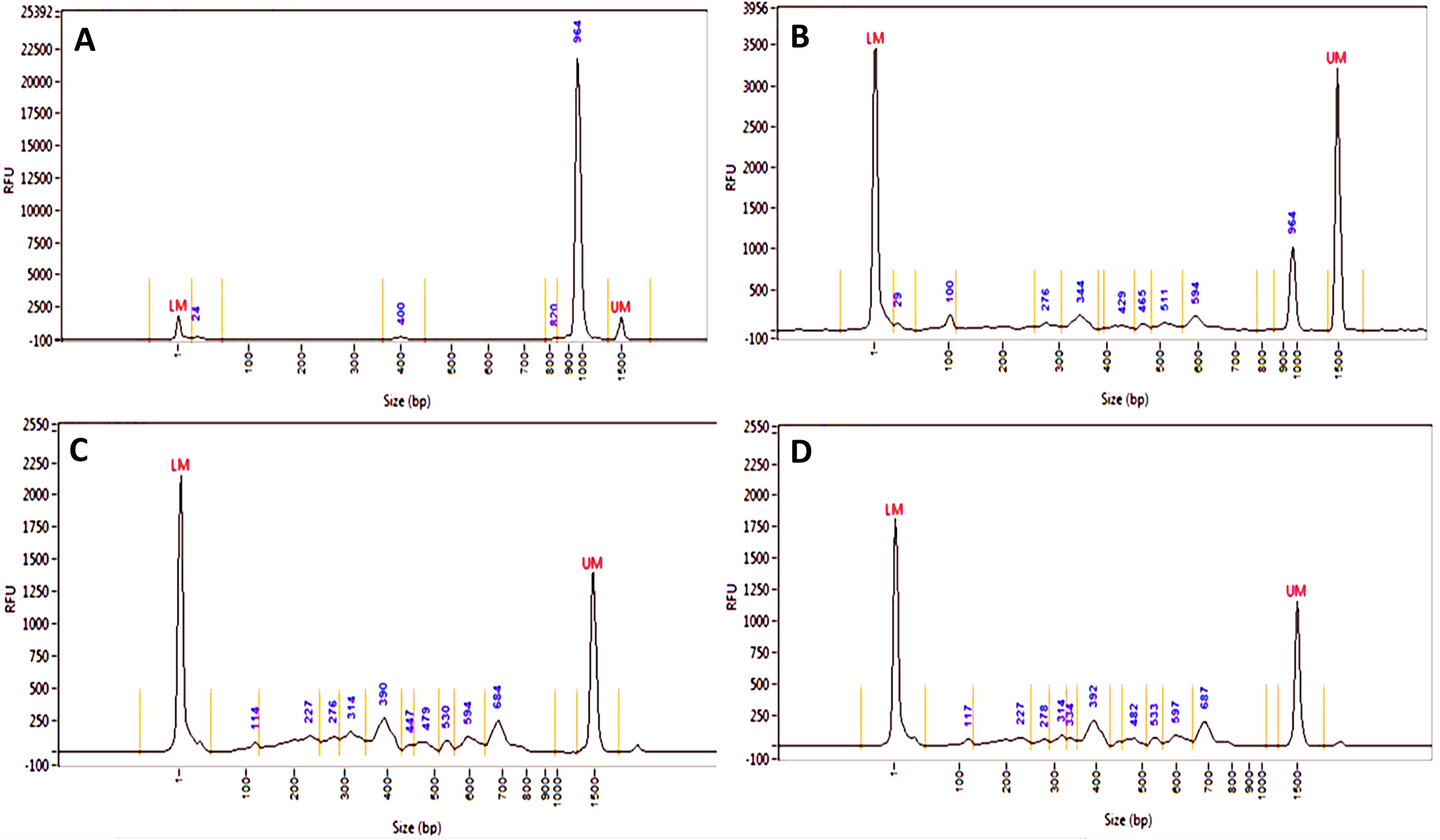
Comparison of different fragmentation parameters of a single PCR amplicon. Panel A shows the non-fragmented control of the single 964 bp PCR product as assayed by capillary electrophoresis. Panel B shows the results of test parameter 1 in Table 1. This is an example of incomplete fragmentation as the original 964bp PCR product can be observed at a high concentration. Panel C and D are technical repeats showing results of fragmentation using test #13 parameters. The y axis in all graphs shows relative abundance of product in relative fluorescence units (RFU). The x axis shows molecular weight in base pairs. LM and UM are the lower and upper marker, respectively.

### Sonication of complex pools of PCR amplicons

Having established optimal conditions for fragmentation of a single amplicon, we next sought to test these parameters in a TILLING by Sequencing experiment containing 32 amplicons ranging between 670-1513 base pairs. PCR primers producing these amplicons had been previously validated and used in traditional TILLING assays employing gel-based cleavage assays using an eight-fold genomic DNA pooling strategy ^12–15^. Thus, the amplicon set represents the complexity of a TILLING project, and also allows testing of the feasibility of applying sonication-based fragmentation and next generation sequencing to an already existing TILLING platform. Fragmentation was performed on 48 pools (see Supplementary Fig. S4) followed by preparation of Illumina libraries. Fragmentation profiles of the prepared libraries showed more complex patterns with average sizes of approximately 650 bp (including index adapter sequences) and the presence of higher concentration peaks at specific molecular weights (Fig. 2).

**Figure 2:**
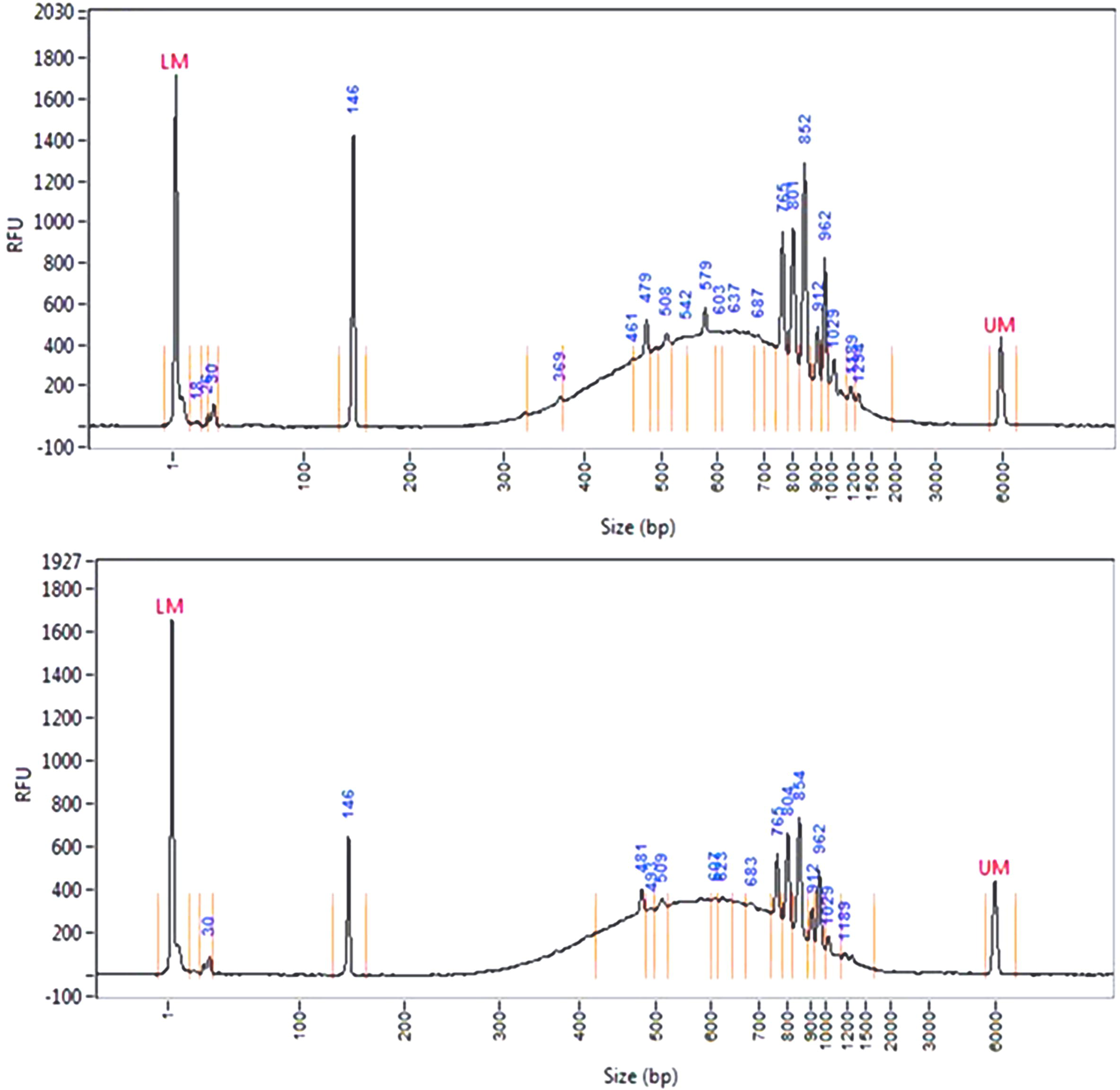
Fragmentation profiles of Illumina sequencing libraries prepared from 32 pooled amplicons produced by PCR amplification of genomic DNA from 256 pooled barley samples. PCR products from forty-eight genomic DNA pools were subjected to sonication using test parameter 13 in Table 1. Fragmentation profiles of two different libraries are shown (top and bottom panel).

### Evaluation of sequencing libraries prepared from pooled fragmented amplicons

To evaluate the effect of fragmentation on sequencing coverage and quality, the Illumina libraries were subjected to 2×300 PE sequencing on a MiSeq instrument. Sequencing coverage profiles, per base mapping quality, and per base sequence quality of four unique amplicons are shown (Fig. 3). Similar data was graphed for two pools and two amplicons from a previously published project where TILLING screening was employed by direct sequencing (without fragmentation) of short amplicons derived from PCR from pools of 64 tomato genomic DNAs^16^ (see Supplementary Fig. S5). To further validate the use of sonication for TILLING assays involving complex pooled PCR products we have screened a subset of the mutant barley pools for sodium azide - MNU mutations and have recovered mutations in two gene targets that were previously identified in the mutant population using traditional TILLING methods^15^ (see Supplementary Table S4). Detailed sequencing statistics for pooled samples used to prepare Fig. 3 and Table S4 can be found as Supplementary Data 1.

**Figure 3:**
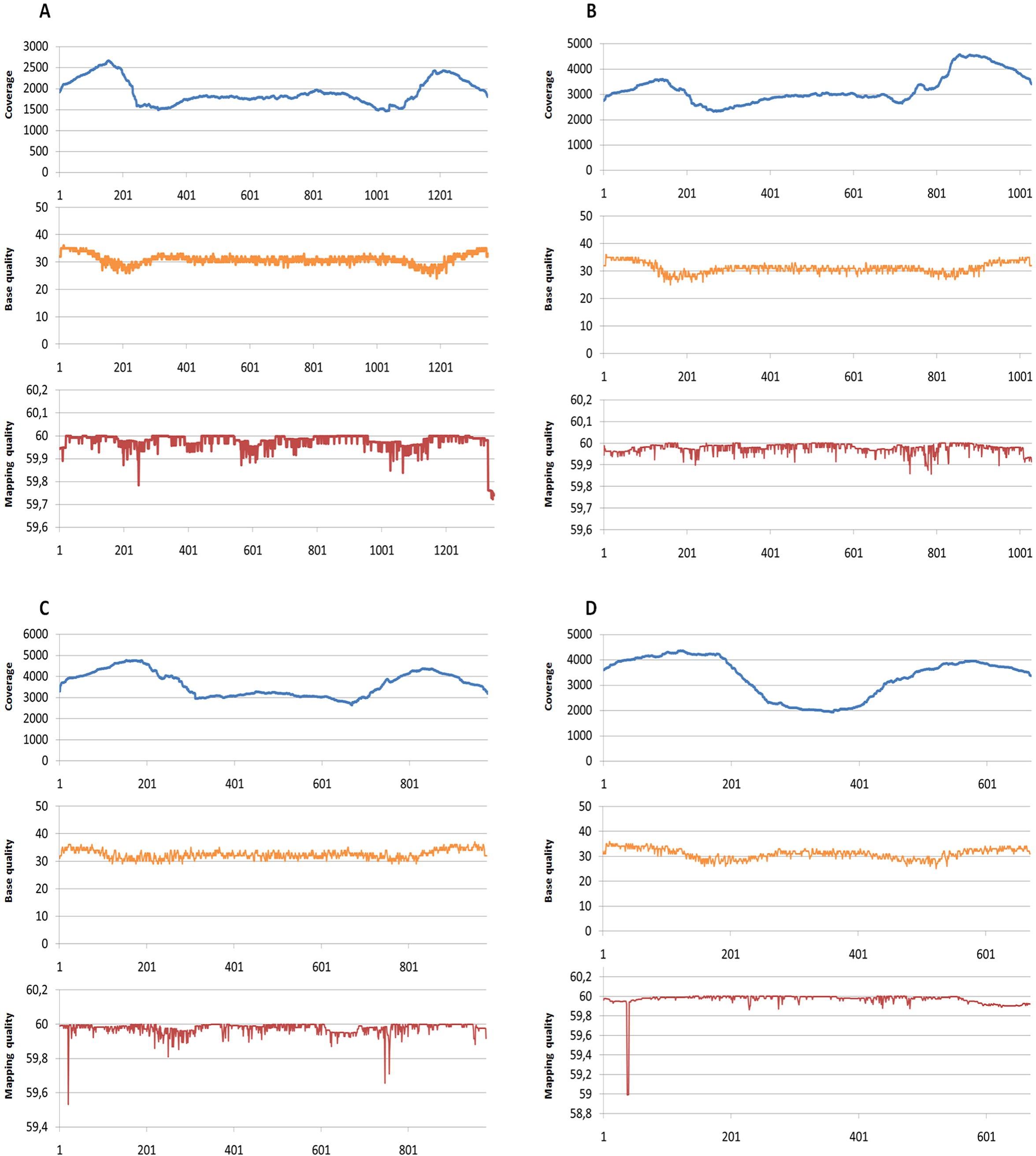
Read coverage, base quality and mapping quality from 2×300PE sequencing of targets EXPB4 (1350 bp) (A), RTH3 (1028 bp) (B), PRT1 (979 bp) (C) and ALS3_1 (670 bp) (D). The number of aligned reads (coverage) at each position in the amplicon is marked in blue. The average base quality per position is marked in yellow and mapping quality in red. Coverage is even across the majority of each amplicon with slight increases in coverage and concomitant decreases in quality score towards the end of each amplicon. Shorter amplicons have longer regions of increased coverage at the ends.

## Discussion

The ability to rapidly discover novel nucleotide variation in germplasm collections and mutant populations is a fundamental tool for functional genomics and breeding of plants and animals. For example, large germplasm collections have been maintained for many species, but limited resources mean little has been done to characterize gene coding and regulatory sequence variation that may prove important for unlocking their full potential.. Further, efficient methods for accurate discovery and cataloguing of nucleotide variation can be important to measure and protect biological diversity in developing countries^17^. Additionally, for functional genomic studies, screening of a TILLING population for the recovery of induced mutations predicted to alter gene function typically requires interrogation of thousands of individuals. While reduced-representation genome methods such as exome capture sequencing have been described to recover rare single nucleotide variants in large genome species such as wheat, the approach remains cost-prohibitive for smaller scale projects and understudied species where research funds are limited^18^. Amplicon sequencing provides a low-cost alternative that is advantageous in that it is highly flexible with regard to population size and the choice of sequence regions for evaluation.

Fragmentation of low molecular weight PCR products is potentially more challenging when compared to working with genomic DNA where fragment sizes can exceed 20,000 bp when using standard extraction protocols^19^. For example, recovery of a 500bp fragment from a 1000bp PCR product requires that a single double strand break be induced at the mid-point of the amplicon. Owing to this, we chose to first seek optimal conditions for fragmentation of a single amplicon. Repeated trials showed that sonication for 30 seconds with a peak power of 50 W, Duty factor of 40 and 200 cycles per burst produced a smooth distribution of fragment sizes that peaked near 400 base pairs, a suitable size for Illumina library preparation and sequencing. While different sized amplicons may require subtle optimizations, we reasoned that sonication at this setting may be suitable for a complex pool of PCR products of varying sizes. We chose to evaluate a pool of amplicons ranging between 670 and 1513 bp because this is the size range of amplicons used in traditional gel-based TILLING approaches^20–24^. We sought conditions to allow higher throughput screening of existing TILLING populations where gene targets and oligonucleotide primers have been previously validated.

To mimic a true TILLING approach, we used large pools of 256 genomic DNAs prepared from a sodium azide - MNU mutagenized TILLING population^15^. We chose 32 amplicons to further mimic a high throughput mutation screen and assayed 48 different pools. When evaluating fragmentation profiles from these pools, a diverse pattern was observed where specific fragment sizes over accumulate. This may indicate that some sequence contexts or fragment sizes are less likely to experience sonication-induced cleavage. This observed pattern, however, did not affect read coverage or mapping quality across the amplicons. It is interesting to note, however, that while read coverage is generally even across fragmented amplicons, there is a consistent pattern of higher coverage and reduced base quality near but not directly at amplicon termini. This pattern is expected in paired end reads if there is a lower probability for fragmentation of DNA sequences near the termini of the PCR amplicon. Importantly, with the exception of the increases near the amplicon termini, read coverage is consistent across the length of amplicons, and sufficiently high coverage was achieved such that mutations can be detected in individual mutant lines represented in the pool. We applied the CAMBa TILLING bioinformatic pipeline for two amplicons and recovered induced mutations previously discovered by traditional gel-based TILLING methods. This suggests that the approach of using high genomic DNA sample pooling and simultaneous discovery of mutations from sonicated PCR products is suitable for the recovery of rare point mutations.

Comparisons between sequencing metrics from fragmented and non-fragmented PCR products allow further evaluation of the sonication-based approach. Data from direct sequencing of non-fragmented PCR products pooled in a similar fashion (see Supplementary Fig. S5) shows both coverage and base quality drop towards the center of amplicons that are near 600 base pairs. Coverage increases in the center of shorter amplicons as expected when applying paired-end sequencing. Based on this data, we conclude that the sonication-based approach produces more consistent data independent of starting amplicon size. This allows for more precise experimental design, taking into consideration genomic DNA sample and amplicon pooling so that required depth of coverage can be obtained for accurate variant calling. Cost estimations can also be considered to further evaluate the utility of sonication of larger amplicons versus direct sequencing. For simplicity, we assume that quality and coverage are identical for a non-fragmented 600 bp amplicon versus a fragmented 1200 bp amplicon, and that all PCR products are quantified using high sensitivity assays on the Fragment Analyzer (FA). Quantification of PCR products is an essential step when pooling multiple amplicons as an equal molar contribution from each amplicon is required to ensure equal representation in the sequencing assay and subsequent data analysis. When considering major consumables, we estimate that it will cost approximately 1.9x more to assay two 600 bp amplicons versus one 1200 bp amplicon that has been sonicated (Table S5). Cost estimations do not take into account the extra time for PCR amplification and FA machine run times when performing more PCR reactions producing shorter amplicon sizes. This adds to the time and cost of assaying shorter amplicons. Additional time and cost savings may be achieved by using amplicons larger than ones reported here. For example, while gene sizes vary, median gene length in plants such as barley can be thousands of base pairs^25,26^. Long PCR, however, can be more challenging to optimize, and careful testing should be performed to avoid unforeseen biases and to ensure that PCR amplification produces and equal representation of molecules from complex pooled genomic DNAs. Errors from PCR must also be considered. Previous studies using traditional gel-based approaches showed using Taq polymerase with amplicon sizes up to 1500 bp resulted in less than a 5% false positive error rate^27,28^. PCR based errors are further mitigated when applying a three dimensional genomic DNA pooling strategy whereby each genomic DNA is represented in three unique pools and true mutations are found only once in each pooling dimension^3^. When applying this criterion to mutation calling the likelihood of false positive errors due to PCR amplification of genomic DNA or through post-ligation library amplification is extremely low. Further, library preparation kits that do not employ post-ligation amplification are available if errors from library preparation are a concern. While in-house sequencing equipment and 2×300PE reads was used for this study, shorter read, higher throughput sequencing platforms can provide greater coverage at lower cost.

We conclude that fragmentation of pooled amplicons by ultrasonication provides a suitable method for producing even coverage sequencing for targeted recovery of nucleotide variation in large populations of samples. Alternative methods for amplicon fragmentation, such as use of dsDNA fragmentase have been applied for TILLING with genomic pooling of 64-fold^29^. This suggests that multiple methods may be considered for experiments employing complex pools of PCR products. Further tests are needed, however, to determine if alternative fragmentation methods will produce similar results in highly pooled samples. In addition to reverse-genetics, we envision that combining sample pooling with amplicon fragmentation can improve screening efficiencies for natural allelic variations. For example, the same approach can be applied for sequencing of germplasm banks where rare alleles may provide an important resource for understanding gene function and allow for the genetic improvement of bred species^5^.

## Methods

### Genomic DNA and PCR

Genomic DNA from *Coffea arabica* was provided by Margit Laimer. PCR primers were designed using the Primer3 program with parameters previously developed for TILLING assays^30^. Primer sequences were TCGATTCGATTCGTTGACACCCCTA and TGGATGATGGATGGGAATGTGGTTC. PCR was performed as previously described^31^. Amplification of a single PCR product was assayed using agarose gel electrophoresis and confirmed via capillary electrophoresis (Fig1.A).

Genomic DNA from chemically mutagenized barley was prepared using a modified CTAB protocol^15^. DNA samples were adjusted to a similar concentration and pooled. Two hundred and fifty-six samples were combined in each pool. Forty-eight unique pools were prepared. Primers were designed to target regions of the H. *vulgare* genome as previously described (Table S3)^32^. PCR was performed on pooled genomic DNA samples using 32 unique primer pairs. Individual PCR reactions were performed for each primer pair. The concentration of each PCR product was then normalized to 200 nanograms (ng) using image J image analysis and a 96-well eGel system^19^. PCR products were then pooled together.

### Sonication of PCR products

Sonication of PCR products was performed using a Corvaris M220 ultrasonicator with microTUBE AFA bead split tubes (coffee amplicons) or microTUBE AFA FibrePre-slit (barley amplicons) in 60 μl volumes except where indicated with parameters adjusted according to Tables 1, S1 and S2. Fragmentation of PCR products was assayed using a Fragment Analyzer with the low sensitivity 1kb separation matrix with 30 cm capillaries (cat #DNF935). Analysis of data was performed using the *PROSize*^®^ Data Analysis Software.

### Sequencing and data analysis

Barley genomic DNA used in this study were previously prepared for traditional TILLING assays^15^. Samples were pooled in a three dimensional array as previously described except that pooling was higher, with 256 unique mutant lines in each dimension^3,16^. PCR amplification using previously validated primers (Table S3) was performed as previously described^30^. Assays were performed in a 96 well plate and PCR products were quantified using a Fragment Analyzer with high sensitivity 1kb separation matrix with 30 cm capillaries (cat #DNF474). All samples were adjusted to a concentration of 200 ng. PCR products were pooled together according to the three dimensional pooling scheme^16^. Sequencing libraries were prepared using TruSeq Nano DNA HT Library Prep Kit (Illumina, Catalogue # FC-121-9010DOC) with 200 ng of starting pooled PCR products according to manufacturer’s recommendations. Sequencing was performed on an Illumina MiSeq using 2×300 PE chemistry according to manufacturer’s protocol. Reads were mapped to amplicon sequences during the sequencing run using the MiSeq BWA default settings ^33^. Sequencing coverage and mapping quality were calculated using Qualimap version v.2.2.1 with default settings except that the number of windows was set to amplicon length to retrieve per base metrics used in Fig. 3^34^. Base qualities were prepared using pysamstats and the baseq feature (https://github.com/alimanfoo/pysamstats). Screening of data for previously identified SNP mutations was performed using the CAMBa pipeline for three dimensionally pooled samples as previously described ^3,16^.

## Abbreviations

bp: base pairs
EMS: ethyl methanesulfonate
FA: Fragment Analyzer
gDNA: genomic DNA
ng: nanogram
NGS: Next Generation Sequencing
PCR: Polymerase Chain Reaction
PE: Paired End
TILLING: Targeted Induced Local Lesions in Genomes

## Declarations

### Availability of data and material

The raw datasets analyzed during the current study are available in the Sequence Read Archive submission #SUB3332618 at https://www.ncbi.nlm.nih.gov/bioproject/422048.

## Authors’ contributions

BJT, JJC, IS & II contributed to experimental design and project oversight. AT, LJ & BJT prepared the manuscript. KG, JJC, MSZ, BJH, & BJT conducted experiments, contributed to data analysis and interpretation and manuscript editing.

## Acknowledgements

We thank Professor Margit Laimer of BOKU for kind donation of *Coffea arabica* genomic DNA. We thank Keji Dada of the Cocoa Research Institute of Nigeria for assistance in coffee primer design and testing

## Authors’ information

ORCID for Bradley J. Till: 0000-0002-1300-8285

